# Tree reconciliation combined with subsampling improves large scale inference of orthologous group hierarchies

**DOI:** 10.1101/417840

**Authors:** Davide Heller, Damian Szklarczyk, Christian von Mering

## Abstract

**Background:** An orthologous group (OG) comprises a set of orthologous and paralogous genes that share a last common ancestor (LCA). OGs are defined with respect to a chosen taxonomic level, which delimits the position of the LCA in time to a specified speciation event. A hierarchy of OGs expands on this notion, connecting more general OGs, distant in time, to more recent, fine-grained OGs, thereby spanning multiple levels of the tree of life. Large scale inference of OG hierarchies with independently computed taxonomic levels can suffer from inconsistencies between successive levels, such as the position in time of a duplication event. This can be due to confounding genetic signal or algorithmic limitations. Importantly, inconsistencies limit the potential use of OGs for functional annotation and third-party applications.

**Results:** Here we present a new methodology to ensure hierarchical consistency of OGs across taxonomic levels. To resolve an inconsistency, we subsample the protein space of the OG members and perform gene tree-species tree reconciliation for each sampling. Differently from previous approaches, by subsampling the protein space, we avoid the notoriously diffcult task of accurately building and reconciling very large phylogenies. We implement the method into a high-throughput pipeline and apply it to the eggNOG database. We use independent protein domain definitions to validate its performance.

**Conclusion:** The presented consistency pipeline shows that, contrary to previous limitations, tree reconciliation can be a useful instrument for the construction of OG hierarchies. The key lies in the combination of sampling smaller trees and aggregating their reconciliations for robustness. Results show comparable or greater performance to previous pipelines. The code is available on Github at: https://github.com/meringlab/og_consistency_pipeline

## Background

From the initial definition of orthology and paralogy by Walter Fitch [1], which distinguishes whether two genes diverged from their last common ancestor by speciation or duplication, the concept has been expanded to the notion of orthologous group (OG) [2]. The latter aims to represent a set of genes from two or more species that are in a homologous relationship with respect to their last common ancestor at a given speciation event. This extends the historically pairwise relationship of orthology to be more inclusive. For example, an OG can contain paralogs, if their duplication occurred after the speciation event of reference. In fact, we distinguish between in-paralogs and out-paralogs when the duplication event occurred respectively after (in) or before (out) the speciation of reference [3].

When defining OGs, one always chooses a taxonomic level of reference, i.e. the last common ancestor of the species included in the OG. Because of this characteristic, many resources have focused their attention on providing hierarchically layered OGs [4, 5, 6, 7, 8] or “OG hierarchies” illustrated in (figure 1). This has proven a useful extension to provide a connection from larger OGs, whose ancestor is distant in time, to more fine-grained OGs, whose species are more closely related [6]. The method used to compute hierarchies of OGs differs across resources. For example, eggNOG [9] and orthoDB [10] compute OGs independently at various radiations on the tree of life, while others rely on a graph based approach [11] or hierarchical pairwise comparison [8].

**Figure 1.**
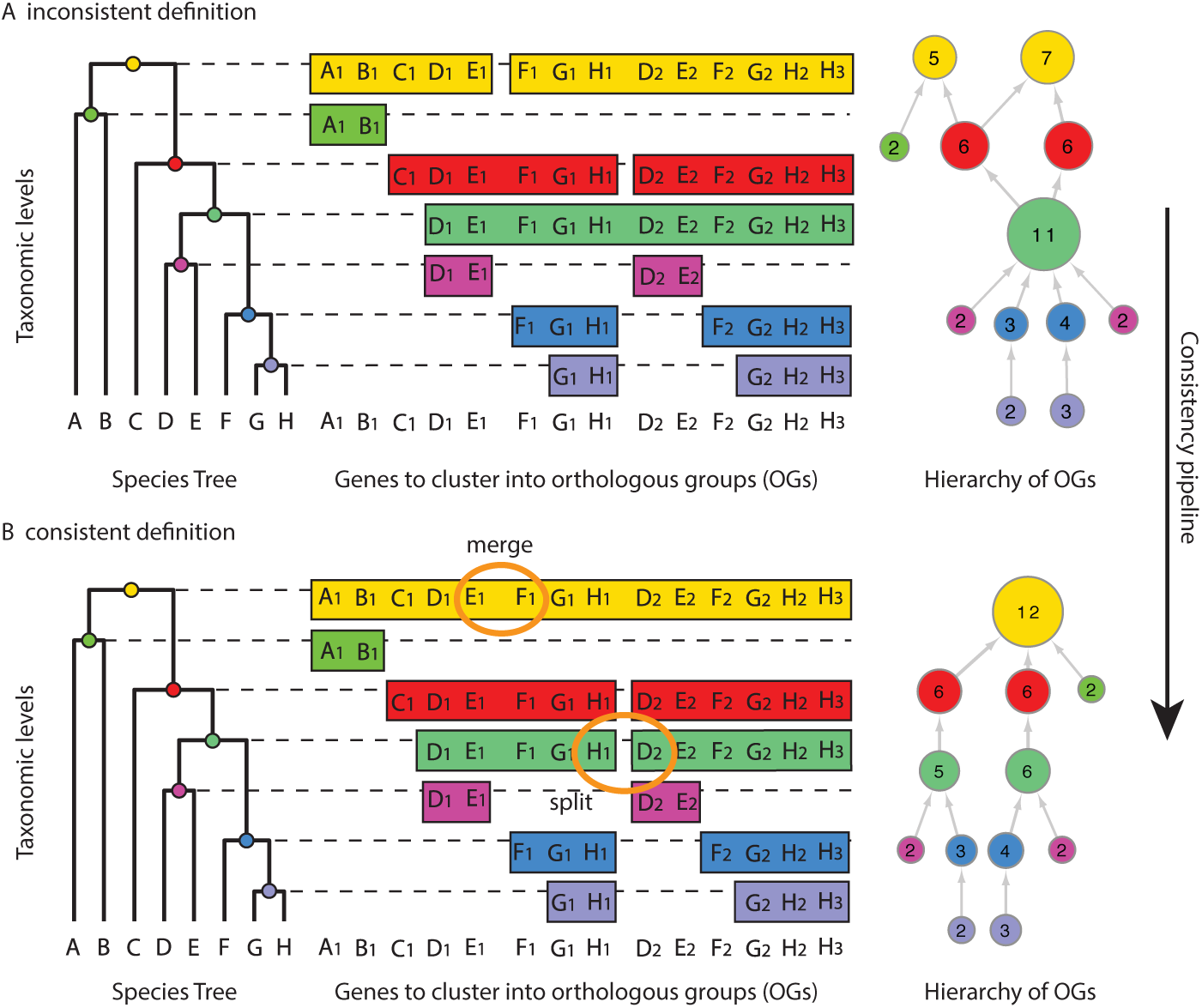
Hierarchical orthologous groups and the consistency problem. The example shows how genes are clustered into orthologous groups (OG) based on the chosen taxonomic level and how the independently computed levels can be joined into a hierarchy of OGs. **(A)** shows a hierarchically inconsistent definition, while **(B)** shows the repaired and consistent definition. Split and merge operations to make the network consistent are highlighted in orange. (Figure based on [12])

Intuitively, when discussing the prediction of OG hierarchies, gene tree inference combined with species tree reconciliation would seem the ideal answer, but it has been difficult to build phylogenies that are sufficiently accurate, while being as computationally scalable as clustering methods [13, 14]. On the other hand, clustering methods, such as eggNOG and orthoDB, must work with varying genomic signal across levels. At every level, the species composition is different and as a consequence the genetic signal will result from different rates of evolution as well as varying quality of genome annotation. It is therefore possible that two independent clustering processes at two different taxonomic levels can create hierarchically inconsistent results (figure 1A). For example, while it would be expected that all the proteins of an OG at the taxonomic level of mammals should be found in a single OG at the vertebrate level, it is possible that the previously clustered proteins split in two separate OGs at the vertebrate level. Such inconsistencies limit the propagation of information across the database and furthermore present the end-user with incompatible results for distinct levels.

Here we present a new methodology to resolve inconsistent hierarchies of OGs based on tree reconciliation. We avoid the problem faced by tree-based methods described above by subsampling the space of proteins that are part of an inconsistency. Sampling small sets of genes from each inconsistency allows to reconcile many phylogenies even for hierarchies containing very large OGs, for which it would be difficult to build accurate phylogenies. Each reconciled tree sample is then used to infer how the OGs should be repaired (figure 1B) to ensure consistency between taxonomic levels. To validate the approach, we have applied the consistency pipeline to the eukaryotic clade of the eggNOG database. We measured the performance through the QFO benchmarking service[15] and InterPro domains [16] and show that by making all OGs hierarchically consistent, we actually improve the performance of the database.

## Methods

The proposed pipeline to resolve inconsistent OG hierarchies consists of six major steps (figure 2): (1) expanding individual OGs to a hierarchical definition connecting several taxonomic levels; (2) sampling the expanded definition by selecting subsets of proteins spanning the inconsistency; (3) building a phylogenetic tree for each of the subsamples; (4) reconciling the sampled gene trees with the species tree using a tree reconciliation algorithm; (5) joining the solutions resulting from the reconciliation to decide how to repair the inconsistency; (6) propagating the applied solution to all lower levels if new inconsistencies are formed. Since the current application of the methodology is the eggNOG database, we will describe the following sections with the latter in mind, but the approach can be adapted easily to other sources. An open source python implementation is available at https://github.com/meringlab/og_consistency_pipeline.

**Figure 2.**
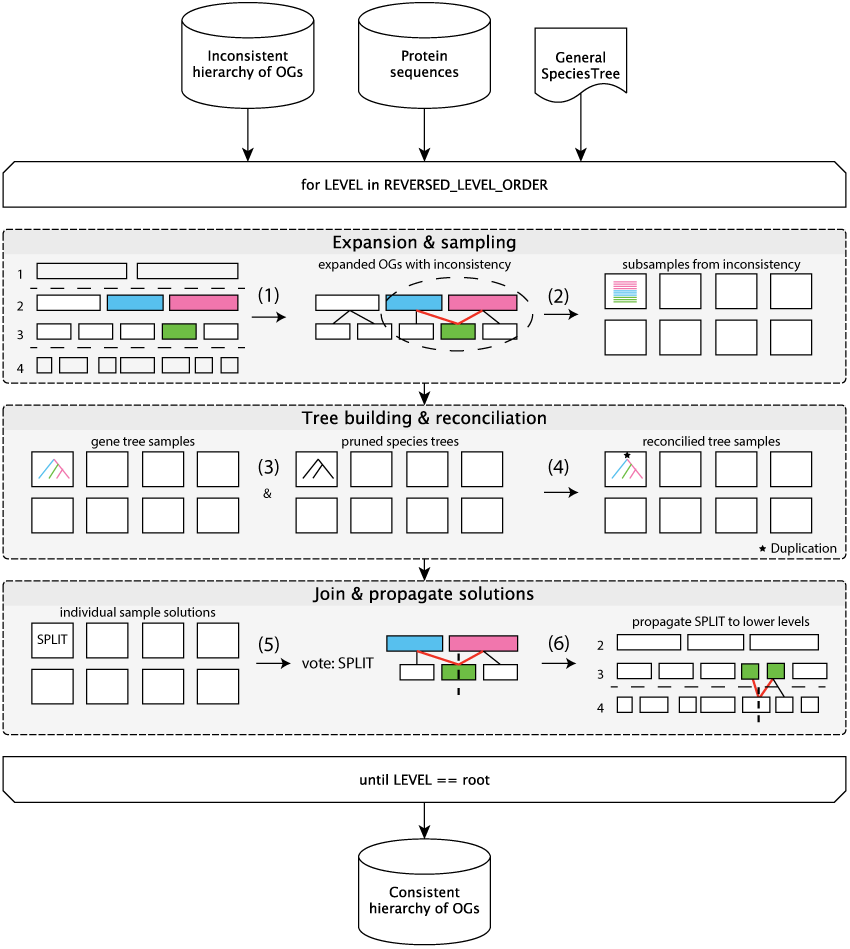
Flowchart of the consistency pipeline. Given an inconsistent hierarchy of OGs, the consistency pipeline traverses the hierarchy of levels in reversed level order, i.e. starting from the leaves, every parent level is visited only after all lower levels have been visited. To make each level hierarchically consistent with all its lower levels, the described six steps are applied: (1) expansion of OGs between one parent level and its children levels, to identify hierarchical inconsistencies (red lines); (2) subsampling of the expanded OG (dashed oval) to obtain sequence samples; (3) Gene tree computation from the sequence samples and pruning of the general species tree to match individual gene tree samples; (4) reconciliation of gene tree and pruned species tree samples; (5) majority vote to determine the solution to resolve the inconsistency, i.e. merge or split; (6) propagation of the split decision if new inconsistencies arise in children’s descendant levels. The algorithm repeats until the root level is completed and the entire hierarchy will be consistent.

### Expansion of orthologous groups

The expansion step consists of detecting hierarchical inconsistencies, by following the protein members of each OG between related levels (parent-children). The parent level is the next higher taxonomic rank, i.e. closer to the root level in the taxonomy tree. For example, in eggNOG’s level hierarchy a higher level would be Supraprimates while its lower levels would be Primates and Rodents. Starting from the proteins of an OG at a lower level, each protein is matched at the higher level to determine whether it is assigned to a higher level OG. For each protein of the matched higher OG, the search process is reversed towards the lower levels, to determine the assignment in the lower levels. The search process continues as long as new OGs are found. In a graph analogy, OGs would be the nodes of a graph where edges represent protein overlap between a higher and a lower OG. In this analogy, the search algorithm simply determines the connected components of the graph. We denote each connected component as expanded OG (figure 2, dashed oval) to represent the fact that it was created by expanding a single initial OG. Hierarchical inconsistencies, are now easily found whenever the proteins of a lower OG diverge in two or more higher OGs. Because of the presence of singletons (single protein member not assigned to any OG), we differentiate inconsistencies when composed of only singletons at the higher level or only one higher OG of size larger than two. These trivial cases are automatically assigned to be merged without further phylogenetic testing.

### Sampling the inconsistencies

To assess via tree reconciliation how to resolve a hierarchical inconsistency, we apply a subsampling strategy. Since OGs can consist of hundreds or even thousands of proteins, it is computationally expensive to build reliable phylogenetic trees including all proteins in the inconsistency. Therefore, we repeatedly sample a subset of proteins and use the latter to build phylogenetic trees for the reconciliation step. The sampling strategy is a guided process, i.e. not entirely random; instead, the species composition should be such that the last common ancestor is located at the higher taxonomic level. This criterion ensures that the tree reconciliation step determines whether, to solve the inconsistency, the higher OGs should be merged (speciation event) or left separated (duplication event) by splitting the lower OGs. In order to fulfill the criterion, the guided sampling process first determines the species composition of all the proteins in the inconsistency. Then, the species composition is used to determine which child taxonomic levels are composing the problem. For example, for an inconsistency at Supraprimates, the species composition could be Primates, Rodents or Leporidae species, added at Supraprimates. By sampling proteins from at least two of these levels, we can ensure that the root of the sampled tree is located at Supraprimates. For the special case in which the species composition comes from only one of the child levels, e.g. primates, a merge decision is automatically assigned without further phylogenetic testing. This follows the assumption, that such inconsistencies are best addressed already at the lower level, whether the inconsistent primate’s proteins should be clustered together or not.

### Tree building

For each sample, we retrieve the protein sequences used to define the orthologous groups (see input data section) and build a phylogenetic tree. The multiple sequence alignment is computed using MAFFT [17] and the phylogenetic tree is built with FastTree [18]. While several other combinations are possible, we chose the latter combination due to its reliability and speed. The resulting trees are rooted with the midpoint criterion, which in absence of reliable outgroup information is a reasonable alternative [19] and is commonly chosen by high-throughput phylogenetic tree workflows [20].

### Tree reconciliation

For every sampled phylogenetic tree, we prune a general species tree until it only includes the species in the sample. The reconciliation between the sampled gene tree and the pruned species tree is performed using the tree reconciliation software NOTUNG [21]. The results of the tree reconciliation predict for each inner tree node which evolutionary event has occurred, using the maximum parsimony principle. We limit the inference to the detection of speciation, duplication (D) and loss events (L), also defined as the DL scenario. While it is possible to include the detection of lateral gene transfer (LGT) events, i.e. the DTL scenario, we do not expect that in the eukaryotic domain of life LGT events have a strong influence on inconsistencies given their limited occurrence [22].

### Solution joining

The evolutionary events resulting from the reconciliation algorithm suggest the solution for solving the inconsistency and making the hierarchy of OGs consistent. Specifically, we focus on the evolutionary event corresponding to the root of the tree to decide whether to split or merge the inconsistency. A duplication event indicates that at the current taxonomic level the two sister clades are in a paralogous relationship and therefore the higher OGs should stay separated. We apply the split decision, to separate the lower, inconsistent, OG into two or more OGs according to the higher OGs by protein overlap. A speciation event indicates instead that the two sister clades are orthologous and should by definition be part of the same orthologous group. We apply the merge decision to join the higher OGs into a single OG. Because for each inconsistency we have multiple tree reconciliations, we separate the join process into two sub-routines. First, individual reconciliation samples are aggregated by majority vote to decide whether to merge the higher OGs or split the lower OG. Second, since the expanded OG can be composed of several inconsistencies, we apply the solutions iteratively until the expanded OG is completely consistent.

### Solution back-propagation

While merge operations change the higher taxonomic level, without influencing the lower levels, the split operations divide OGs at the lower levels, making it possible that new inconsistencies arise in the subtree of the descendant levels, which was previously consistent. Using the eggNOG level hierarchy as example, if an inconsistency between the higher level Mammalia and the lower level Superprimates is solved by splitting the OG at the Superprimates level, this may create an inconsistency with its descendant levels, Primates and Rodents. To ensure maintenance of consistency, split operations are therefore back-propagated towards the leaf levels whenever new inconsistencies arise, that is, the conflicting OGs in lower levels of the hierarchy are split as well until no inconsistency is present in the sub-hierarchy.

### Methods and data for validation

#### Input data

We applied the consistency pipeline to version 4.0 of the eggNOG database [23], which provides nonsupervised orthologous groups for 2031 species across 107 taxonomic levels. For this study, we focused on repairing the eukaryotic clade of the eggNOG level tree containing 238 species and 2’859’900 clustered proteins. In this particular subset of the database there were 273’784 inconsistencies, out of which 63’846 classified as non-trivial (see sampling method). The pipeline was applied to each level in the hierarchy in reversed level order (leaf to root), such that for every level the children are either leaves or have already been repaired to be consistent towards lower levels.

#### Species Tree

The species trees used for reconciling gene trees in the reconciliation step are pruned versions of the same general species tree. The latter species tree was computed using 40 marker genes and the NCBI reference taxonomy [23].

#### Algorithm parameters and performance

The consistency pipeline has two parameters that determine the sampling algorithm, the number of samples (n) and the number of protein sequences in each sample (m). For the results shown below, we used n=30 and m=25, leading to a total average of 1’214’808 reconciled trees to resolve all non-trivial inconsistencies. The tree computations and reconciliations were parallelized on an SGE cluster with 600 cores. The remaining tasks, including expansion, sampling and joining were performed on a high memory machine using 10 cores. Overall, the execution of the pipeline lasted on average 21 hours (of which cluster operations: 10h tree computations, 3.6h reconciliation).

#### Third party software parameters

MAFFT (v6.861)[17] was used with default parameters (–auto) and –memsave when the input sequence was above 4000 amino acids; FastTree (v2.1.9)[18] with -nopr -pseudo -mlacc 3 -slownni options for increased reconstruction accuracy; NOTUNG (v2.9)[24] with default weighing scheme (D=1.5, L=1) and the – rearrange option with threshold 0.9. The latter option rearranges the topology of the tree for weakly supported branches to minimize the cumulative cost of the reconciliation.

#### Quest for Orthologs (QFO) benchmark

We used the orthology benchmark service published by Altenhoff et al. [15], which offers a series of benchmarks for the evaluation of orthologous gene pairs, using 66 reference proteomes, to provide a “common denominator” for method comparison in the orthology field. We continue to support this effort but acknowledge, together with the benchmark authors [15], that the included tests are geared towards orthologous pair predictions and less optimized for OG predictions and, to an even lesser degree, for hierarchical OG definitions. Furthermore, given the reduced number of reference proteomes, it was not possible to define a large taxonomic hierarchy for which to apply the consistency pipeline. We therefore mapped the QFO proteomes through the reciprocal best hit method to the eggNOG proteomes and designed a simple bottom up algorithm to convert the hierarchy of OGs into pairs of orthologs (see S.I.). Given the high degree of variability introduced by such conversion process, we used the benchmark service as a control to test whether the orthologous pair prediction performance changed after applying consistency rather than to compare against other methods submitted to the QFO benchmark.

#### Domain benchmark

We devised a benchmark that is better suited for larger number of species and hierarchies of OGs, by analyzing the protein domain distribution across OGs. From a structural point of view, domains constitute the functional units of proteins but at the same time, from an evolutionary perspective, they are also highly conserved protein sequences [25, 26]. They are the most straight forward building blocks of deep homology [27] and as such tightly connected to the definition of OGs. As originally shown by [2], OGs tend to closely represent conserved domain families. The assumptions for this benchmark are twofold: in a hierarchy of OGs, the OG at the taxonomic level closest to the evolutionary origin of the domain, should (1) contain all annotated proteins and (2) exclude the proteins with conflicting domain annotation. Since these assumptions are not without challenges [26], we have excluded domains with short sequence length (*<*= 50aa) given their high degree of mobility [28, 29] and excluded potential cases of convergent evolution (e.g. zinc-fingers and repeats). Furthermore, since we applied our pipeline to the eukaryotic levels of eggNOG, we restricted the benchmark to domains that were annotated exclusively in Eukaryotes and present in at least 5 species.

To annotate the proteins in the eggNOG database we used the domain database InterPro(v64.0) as well as the UniProt database to link protein identifiers. We restricted the test set of eggNOG proteins to a subset with high confidence matching to UniProt (1- to-1 mutual best hits and above 70% sequence identity). This condition limits the maximum number of available annotations, but minimizes the error rate of incorrect-annotation due erroneous mapping. We use the mapping file available from InterPro FTP service (protein2interpro.dat.gz, version 64, [16]) to identify the UniProt ids to annotate. Additionally, we selected the domains originating from the PFAM database to define a consistent source of annotation and further pruned the dataset as described above. The final set of tested domains can be further divided into InterPro Domains and InterPro Families [30]. The total number of tested domains is 4120, out of which 2143 are defined as InterPro Family and 1977 as InterPro Domain.

For each domain, we matched all the orthologous groups at every taxonomic level that contain at least one protein annotated with the tested domain, i.e. at least one true positive (TP). We considered as false positives (FP) only proteins from species annotated with the tested domain (or positive). Therefore, the benchmark does not account for species that entirely lack annotations of the tested domain. We also excluded from testing proteins that were not matched with sufficiently high score to UniProt. We computed precision (or positive predictive value, *PPV* = *TP/*(*TP* + *FP*)), recall (or true positive rate, *TPR* = *TP/*(*TP* + *FN*)) and their harmonic mean or F1 score, to be used as the final score with which to validate the benchmark. For each domain, the best F1 score is selected across all levels.

#### Ad-hoc consistency strategy for eggNOG 4.5

We have developed a temporary method of forcing the consistent hierarchy by repairing the obvious clustering errors without introducing significant deviation from the size distribution of the original OGs[9]. As with the presented reconciliation-based method, the algorithm is applied in reverse (leaf to root) level order prioritizing the human branch. For every inconsistency between the tested and the parental level, the method is applied iteratively between the largest parental cluster and every other affected parental cluster in the descending size order. The decision consists of either joining the parental clusters (merge) or splitting the proteins from the tested cluster (split) at the tested level and every affected branch in the leaf-ward direction. The method, by default, joins the inconsistent clusters when there is no overlap between species. If such overlap exist the merge decision is applied only to groups with dissimilar sizes (more than 100% size difference) otherwise the split decision is applied. Such blunt heuristic seems not to create very large Ogs which are not present in the original clustering.

## Results

We present the benchmark results by comparing the consistent OG definition generated by our method (Level Sampling LS) against OG definitions generated, using the same eggNOG dataset, by the following methods: the original inconsistent version (v4.0), the ad-hoc strategy for eggNOG v4.5 (v4.5), a strategy that always merges inconsistencies (MERGE – M), always splits inconsistencies (SPLIT – S) and randomly chooses whether to split or merge an inconsistency with equal probability (RANDOM – RND). Besides the original definition (v4.0) all other OG definitions are hierarchically consistent.

In (figure S4) we show the results from the QFO benchmark service. Since the presented method (LS) consists solely of eukaryotic OGs, the ortholog predictions from missing species pairs are taken from (v4.0), the baseline. The results showed close performance between the baseline (v4.0), the presented method (LS) and the ad-hoc strategy (v4.5), while the (SPLIT) scenario departs from the baseline (v4.0) with a much lower number of predicted pairs (lower recall). It was not possible to identify a clear winner across the benchmark, with all versions often aligning orthogonally to the direction of best performance. For general test categories, on average, the presented method (LS) shows better performance in the species tree discordance tests (fig S4A,B), while the ad-hoc strategy (v4.5) a better recall and precision in the reference gene trees tests (fig S4C,D). RND and MERGE were not included due to the high number of generated orthologous pairs (respectively more than 8 billion pairs for MERGE and 3 billion pairs for RND, compared to less than 21 million pairs for each of the remaining methods).

In (figure 3) we introduce the results of the domain benchmark with the example domain Pop3 (InterPro Family IPR013241: RNase P, subunit Pop3). This domain is annotated in 47 fungal species, with each 1 annotated protein. The line profiles connect the best matching OG for each version for a hierarchy of nested taxonomic levels, in this example from Saccharomyceta over Ascomycota, Fungi, Opistokonta to Eukaryota. The inconsistent version (v4.0) represents our baseline performance. The line profile shows that the best scoring OGs do not include all annotated proteins (17 at most) with the consequent low F1 score. This is also visualized in (figure 4A), which shows the OG network as explored by the expansion step (see methods) before repairing inconsistencies. Inconsistencies are the branching points in the upwards direction and reflect the reassignment of proteins. The random strategy, albeit consistent (figure S1), did not improve the performance of the OG definition by resolving the inconsistencies. The ad-hoc strategy based on species-overlap, performed better but still missed the majority of annotated proteins. In this example, our method (LS) obtained the best score by merging the inconsistencies into a single OG (figure 4B).

**Figure 3.**
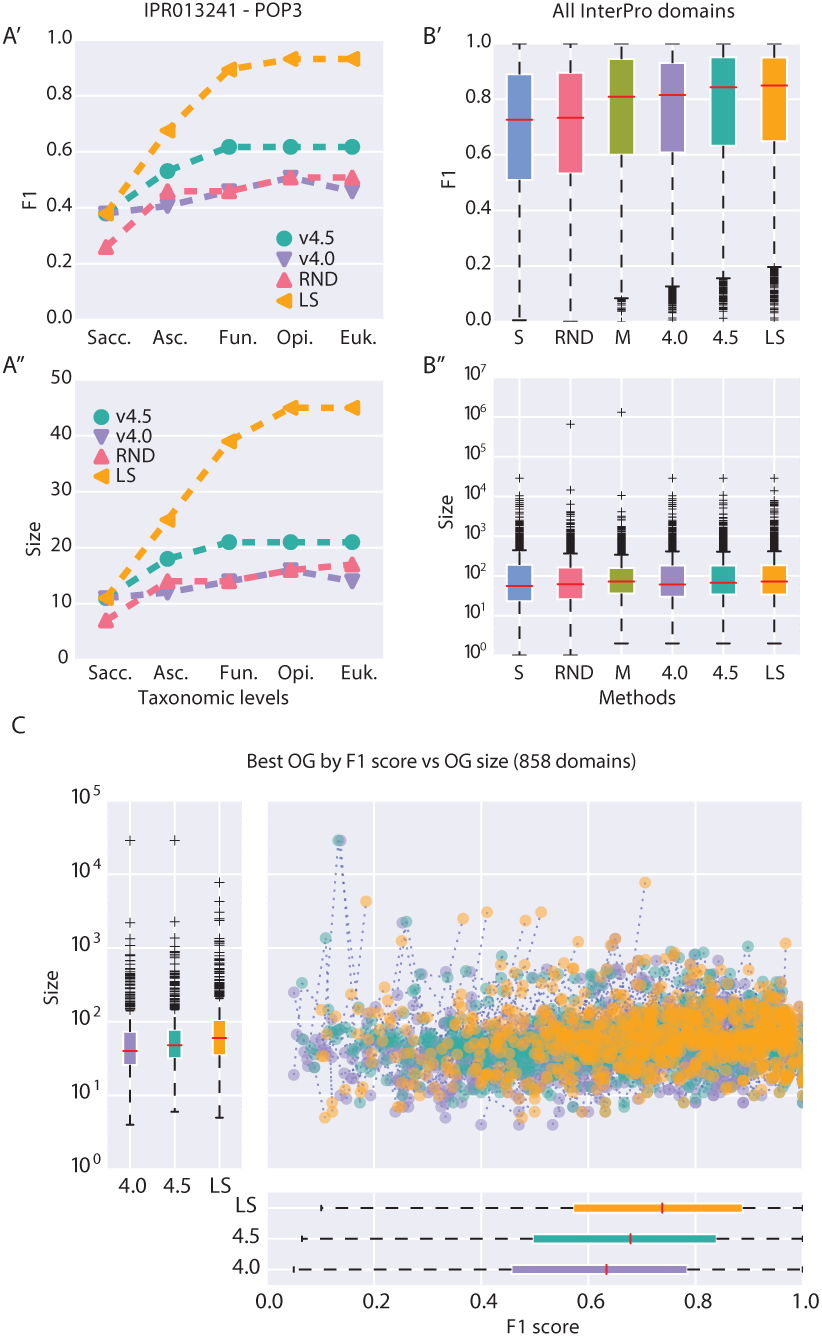
**Domain benchmark results for** tested OG definitions (S/SPLIT; M/MERGE; RND/RANDOM; Level Sampling/LS; 4.0/v4.0; 4.5/v4.5).(**A’-A”**) Example evaluation of InterPro entry IPR013241 (RNase P, subunit Pop3). F1 score and size (no. of proteins) are shown for the orthologous group (OG) with the best matching (F1) to annotated proteins (n=47), for each tested definition (colors) and selected taxonomic levels. Levels in nested order: Saccharomycetales, Ascomycota, Fungi, Opistokonta, Eukaryota. **(B’-B”)** Cumulative results for all tested domains (n=4120). For every OG definition (columns), for each domain, the best matching OG (F1 score) across all tested taxonomic levels is chosen. One-sided paired Wilcoxon signed rank test, alternative hypothesis F1(v4.0) F1(LS) *<* 0, p-value: *<* 0.0001 (all cases); F1(v4.5) - F1(LS) *<* 0, p-value all domains: 0.06 (InterPro Family type 0.02 (figure S2A’) and non-significant for InterPro Domain type (figure S2A”)). **(C)** Selective comparison on 858 domains that differ more than 0.1 in F1 score between the compared methods (LS, v4.0, v4.5). Every point in the scatterplot represents F1 score and size of the best matching OG.

**Figure 4.**
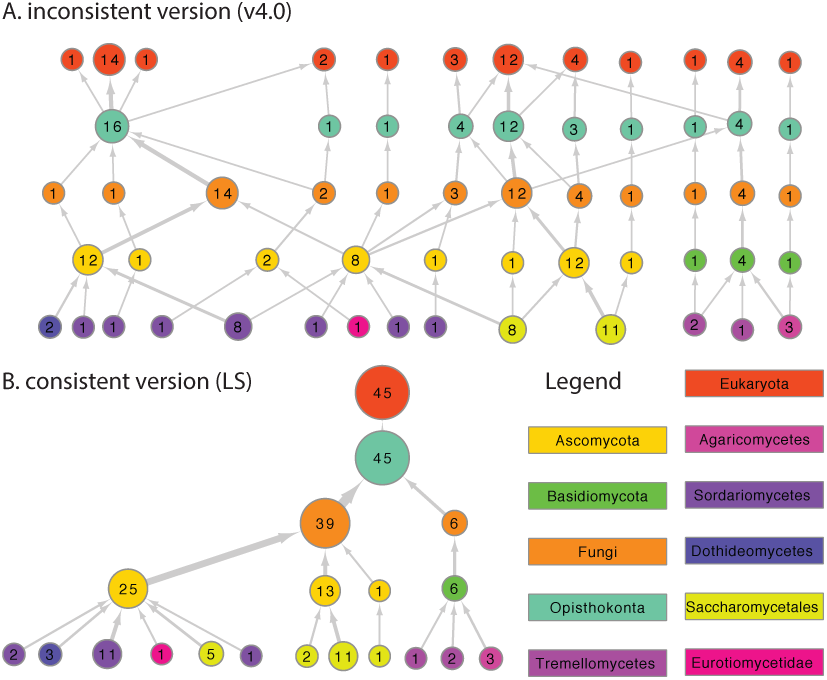
**Expanded OG network example** for proteins annotated with InterPro entry IPR013241. Circles represent individual OG scaled for size and connected to OGs at different taxonomic levels (legend) to represent protein overlap. Labels within the circle represent the size of the OG corrected for proteins which were not mapped to UniProt (see data methods). The network represented in **(B)** shows the same network as in **(A)** after application of the consistency pipeline. Both networks were pruned to the levels represented in figure 3A, for complete networks see figure S1.

To combine findings for individual domains, we selected the OG with the highest F1 score among all levels as representative for the respective OG definition. The boxplots in (figure 3B) show the aggregated results for 4120 InterPro domains. The always split strategy obtained the lowest performance and generates a smaller OG size distribution (figure S2C). Similar performance is obtained by the random strategy, which created both smaller and larger aggregations. The always merge strategy obtained performance similar or better than the inconsistent version, but also generates a very skewed OG size distribution, culminating in a very large cluster at the Eukaryota level (1’533’579 proteins in one OG out of 2’859’900 cumulatively clustered level proteins, 53.62%). The presented method (LS) performed better than the inconsistent version (v4.0) in all tested scenarios (one-sided paired Wilcoxon signed-rank test, p-value *<* 0.0001) and has comparable or better performance than the ad-hoc strategy (v4.5) (one-sided paired Wilcoxon signedrank test, p-value for all domains 0.06 (figure 3B’), for InterPro Family type 0.02 (figure S2A”) and nonsignificant for InterPro Domain type (figure 3A’)). We simplified the difference for visualization in (figure 3C) by representing only the domains for which the F1 score difference between the methods (LS, v4.0, v4.5) is greater than 0.1. Importantly, the presented LS method did not alter the OG size distribution extensively.

## Discussion

In the presented methodology, inconsistencies are solved using one of two possible decisions: (1) splitting the lower OG by subdividing it according to intersection with the OGs at the higher level, resulting in two or more smaller orthologous groups; (2) merging the higher OGs into one single OG, containing all proteins from the OG at the lower level. From this perspective, the reconciliation strategy is one of many possible binary strategies that indicates for every inconsistency whether to split or merge. The resulting OG definition is consistent as long as the chosen strategy does not create new inconsistencies, for example by back-propagating split decisions (see methods).

Notably, also choosing randomly whether to split or merge an inconsistency with back-propagation fulfill the requirements to obtain a hierarchically consistent definition. While consistent, the results for a random or constant decision (always merge/split) were not favorable, and led to either low performance and/or excessive aggregation. A very large and unspecific orthologous group is in contrast with the aims of the OG prediction which seeks to represent groups which can be described as protein families with a common functional characteristics [2]. Indeed, in the introductory example (figure 3A) we see both F1 score and OG size (number of protein members) plateau for the last two levels. This is expected, as the sequence similarity is potentially too small to produce larger clusters at the Eukaryota level, despite speculations that the InterPro Pop3 domain is likely to be related to human RNase P subunit Rpp38 ([31], InterPro description for IPR013241). In the same example, the InterPro annotation is also not represented entirely by a single OG, which is due to the fact that all tested methods rely on the initial clustering (v4.0). More precisely, if in the original definition the OGs containing all annotated proteins are never linked across taxonomic levels by an inconsistently classified protein, the methods targeted at repairing inconsistencies, will never be able to form a single OG containing them (figure S1 v4.0). While it is possible to develop methods that aim to change also hierarchically consistent OGs, it is beyond the scope of the current method to build entirely new OG definitions. On a related note, given that the example annotation is strictly contained in the Fungi kingdom, it would be correct to assume that the best F1 score should be already found at the Fungi level and not at Opisthokonta (figure 3A’). This can be understood considering that the presented methods only merges OGs in the upwards direction, that is, with respect to the OG at the lower level, which proteins are divided into several OGs at the next higher level. The decision of merging lower OGs is possible but does not resolve the inconsistency. On the contrary, the aggregation of lower OGs can potentially create new inconsistencies.

### QFO benchmark

The close clustering of the presented method (LS) to the ad-hoc strategy (v4.5) and to the baseline (v4.0) in the QFO benchmark results (fig S4) is a general indication, that the consistency pipelines preserve the pairwise orthologs prediction performance, but it may also indicate that the tests are not sensitive enough for testing differences in hierarchical OGs. The high recall and precision of the ad-hoc strategy (v4.5) method in the reference tree tests can be partially explained by the more conservative approach in merging OGs, which artificially limits the maximum size of OGs. The presented version (LS) of our method, does not include such size limiting heuristic. We have implemented similar strategies in the production version for the next release of eggNOG (v5.0), bound by computational resources given the high number of included species (5090). This version obtained better recall and precision scores by adding pre-processing operations for OG size control, suggesting that such simple heuristics can indeed increase the robustness of a consistency method when the signal to noise ratio is low. Similar performance improvements can be expected if more computational resources were allocated for more extensive sampling, improved gene tree prediction or tree reconciliation.

### Method parameter space

Overall the results of the proposed consistency pipeline show comparable or better performance (figure 3C, S2) than current methods, suggesting that sampling the protein space of the inconsistencies and using tree reconciliation can correctly determine how to solve an inconsistency. The method does, however, have a large parameter space. Being composed of several individually challenging parts, such as the phylogenetic tree construction and reconciliation, it is virtually impossible to do an extensive parameter screen. Phylogenetic workflows offer a plethora of possible tool and parameter combinations and can vary greatly in computational demand and accuracy. Tree reconciliation in addition, proposes a large number of inferable evolutionary hypotheses and requires individual event cost parametrization in the maximum parsimony framework [32]. For these reason, we guided our choice by practicality (rather than result optimization) and selected a combination of established methods (MAFFT[17], FastTree[18], NOTUNG[24]) for a consistent baseline performance and high throughput capability.

Another important set of parameters of the consistency pipeline govern the sub-sampling step. Sample size (M) and sample number (N) determine the size (no. of sequences) and the number of reconciled trees per inconsistency. To choose the used combination for the results (LS), we performed a convergence analysis of 208 large OGs at the Bilateria level with a starting size (before expansion) of at least 50 proteins. We measured convergence by computing the ratio of the reconciliation outcome between inferred duplication over speciation events (D/S ratio). In (figure S3A) we show that for each choice of M the D/S-ratio converges with increasing N. In most cases (155/208, 73%) the final convergence values were in close vicinity (figure S3A’A”’, std *<* 0.15) while the remaining had larger divergence (figure S3A””). We explain this divergence, by the fact that a smaller sample size inherently limits the maximum amount of sequence variation captured by the tree sample. While it is computationally infeasible to always sample large trees, we chose a value of M=25 that reduces the variation of the D/S scores towards higher M values (figure S3B), while still being computationally feasible. To choose the sample number N, we computed confidence intervals (0.95) along the convergence by comparing the outcome to a Bernoulli process with unknown success probability and correcting the estimate for small sample size [33]. The same results were also confirmed by the standard Wald method[34]. As one might expect the confidence interval initially widens to then decrease along N (figure S3C). We chose N=30 to combine the computational limitations and the reduction in variance with increasing N (figure S3D).

The sample composition is also important for the final step of the reconciliation algorithm, as the merge-split decision is taken based on the evolutionary event inferred at the root of the tree. While we include multiple branches of life by selectively sampling the protein space (see methods), tree rooting remains a difficult phylogenetic problem. A possible improvement for this could be the selection an outlier sequence in each sample to make the inference of the root easier. Another promising improvement lies in a recently proposed aggregated tree reconciliation or super-tree reconciliation[35], holding the possibility of more accurate results.

## Conclusion

We developed a new methodology to solve inconsistencies in hierarchies of OGs based on tree sampling and gene-species tree reconciliation. Against previous results, suggesting that tree reconciliation has limited applicability in the context of large scale inference of OGs [14, 13], we show that it can be effectively employed to determine the solution to hierarchical inconsistencies if combined with sub-sampling. We apply the consistency pipeline to the eggNOG database in the eukaryotic domain of life and evaluate results using protein domain annotations from the InterPro database. Benchmark results reveal comparable or greater performance than the original inconsistent version or the currently employed ad-hoc consistency strategy. Based on these results we have successfully used a slightly modified version of the methodology for the upcoming release (v5.0) of the eggNOG database, with 5090 genomes and 351 taxonomic levels. Despite computational demands limiting the exploration of the large parameter space governing the pipeline, the approach offers several points to further improve the performance. The sampling strategy for inconsistencies could be improved by take into account genome quality measures, such as BUSCO by Simão et al. [36], or known variations in the speed of genome evolution, favoring slow evolving genomes over fast evolving ones. The field of tree reconciliation has seen exciting new theoretical advances which expand the number of modelled evolutionary events, reconciliation for multi-domain families [37][38] or new ways to aggregate results from multiple reconciliations [35]. These improvements can both strengthen and expand the capabilities of our pipeline, making it also suitable for applications in Bacteria and Archean domain as well.

## Author’s contributions

DH, DS and CVM designed the research, analyzed the data, and wrote the article. DH implemented the method.

## ACKNOWLEDGEMENTS

The authors wish to thank Jaime Huerta-Cepas, for helpful discussions and help in using the ete tookit [20], and João F.Matias Rodrigues for helpful discussions and technical help in implementing the program’s execution on the computational cluster.

## Supplementary Figures

**Figure S1.**
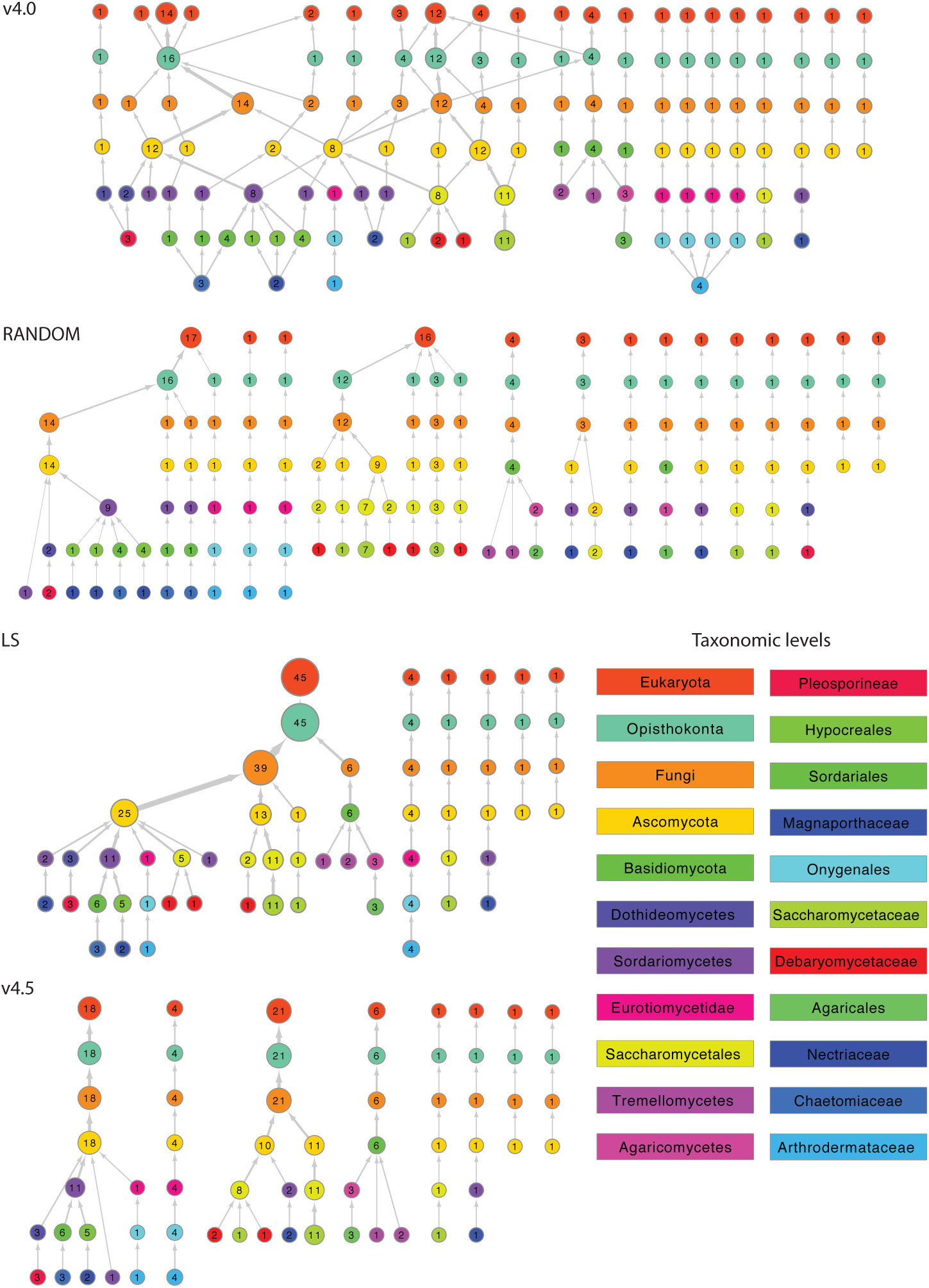
**Fully expanded OG networks** for proteins annotated with InterPro entry IPR013241. Circles represent individual OG, scaled for size (no. of proteins, also shown as label), connected to represent protein overlap to OG at different taxonomic levels (legend). Version 4.0 shows the original OG network with hierarchical inconsistencies while the other methods (RANDOM, v4.5, LS) show the repaired and hierarchically consistent OGs.

**Figure S2.**
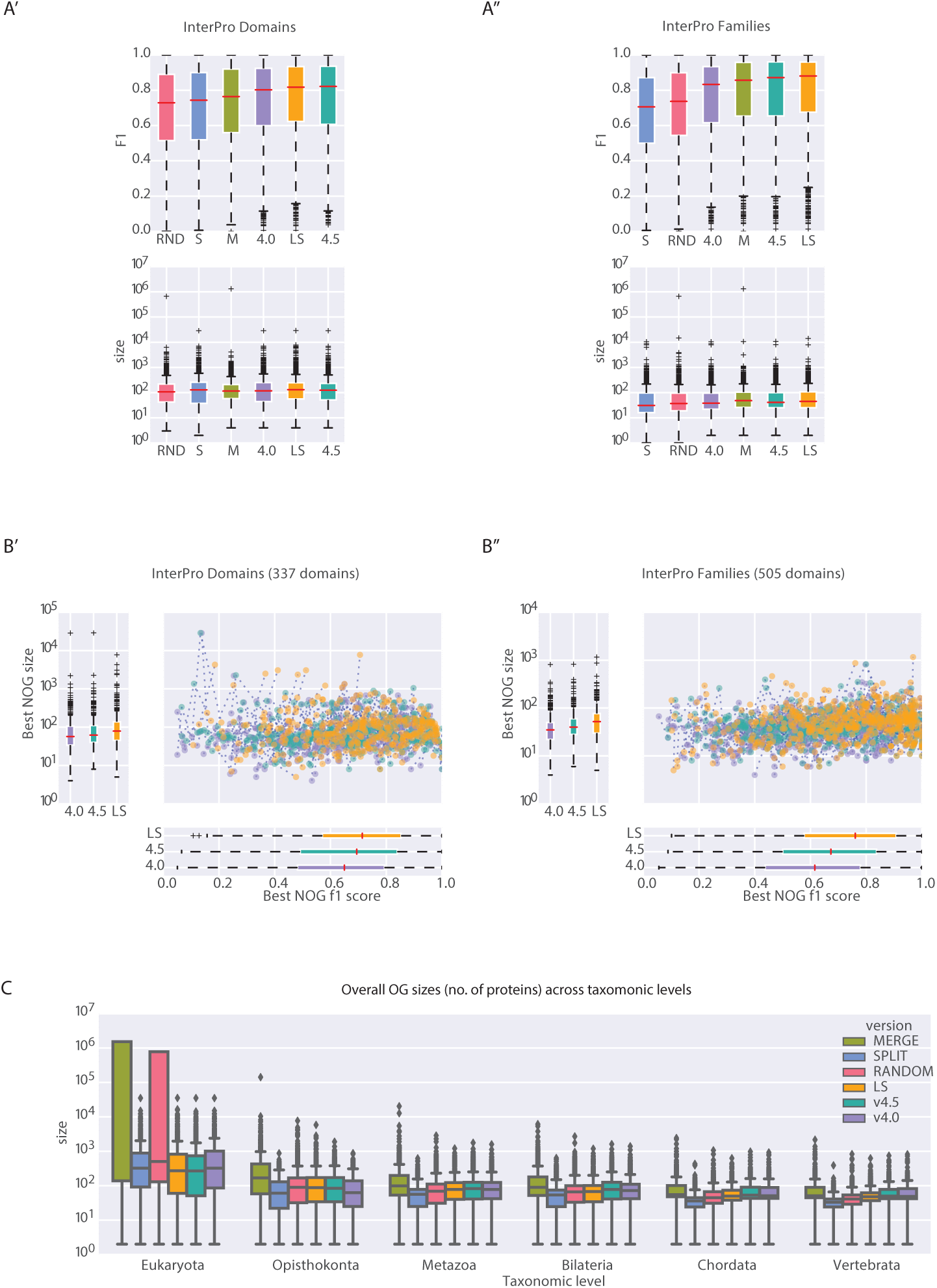
**Domain benchmark results** for sub-types of InterPro annotation and overall OG sizes. **(A’-A”)** Cumulative results for separated domain sub-types, InterPro Domain type (n=1977) and InterPro Family type (n=2143). For every OG definition (columns), for each domain, the best matching OG (F1 score) across all tested taxonomic levels is chosen. One-sided paired Wilcoxon signed rank test, alternative hypothesis F1(v4.0) - F1(LS) *<* 0, p-value *<* 0.0001; F1(v4.5) - F1(LS) *<* 0 for InterPro Family type 0.02 (A”) and non-significant for InterPro Domain type (A’). **(B’-B”)** Selective comparison on domains that differ more than 0.1 in F1 score between the compared methods (LS, v4.0, v4.5). Every point in the scatterplot represents the F1 score and size of the best matching OG. Subdivided by InterPro domain type: Domains (n=337) and Family (n=505). **(C)** Overall distribution of OG size (no. of proteins) per version and taxonomic level.

**Figure S3.**
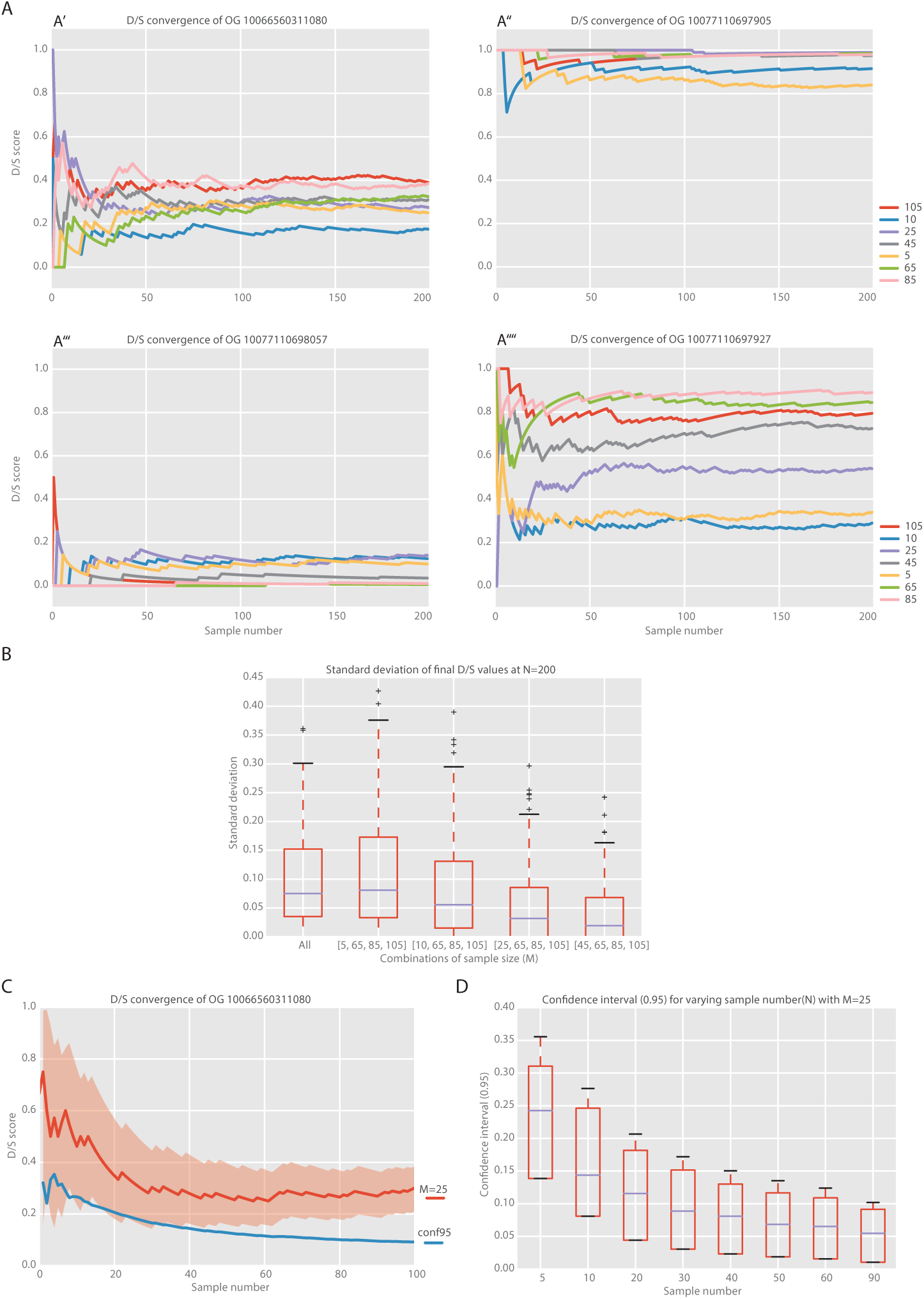
Convergence study for sampling parameters. **(A)** Duplication-Speciation ratio (D/S) convergence for selected sample sizes (M = 5,10,25,45,65,85,105) over 200 samplings (N). A’-A”’ show examples of solution convergences with low variation across M; A”” shows an example with large variation. **(B)** Quantification of D/S variation across all tested OGs (n=208) at N=200 for different M combinations. The first including all M values and the other three large values (65,85,105) combined with one smaller value (5,10,25,45). **(C)** Confidence quantification for D/S of M=25 across increasing N values for example A’ using an exact estimation for binomial processes [33]. **(D)** Quantification of confidence interval size over all tested OGs (n=208) for M=25 across increasing N values.

**Figure S4.**
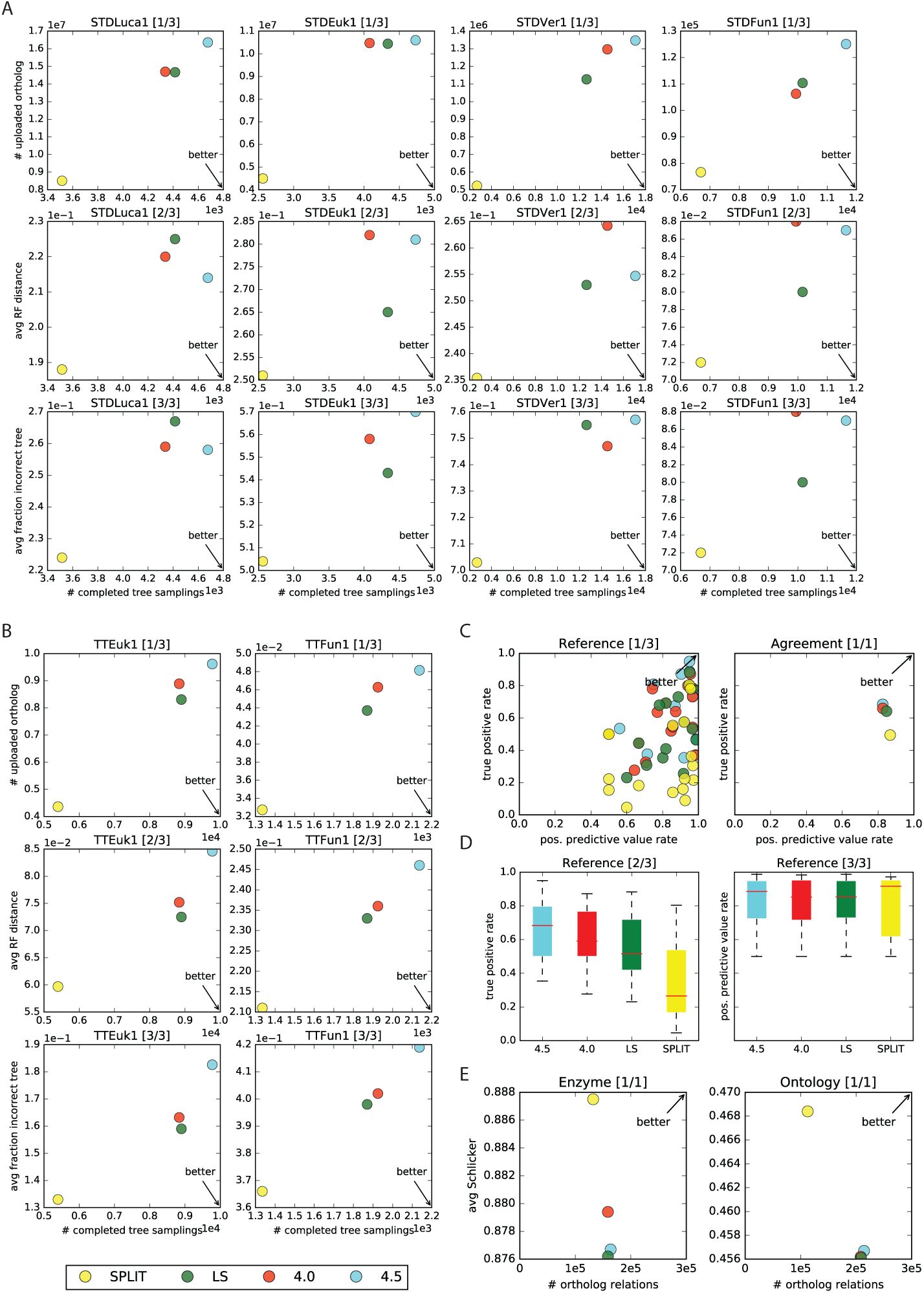
Quest for Orthologs Benchmark. **(A)** Generalized Species Tree Discordance Benchmark (Altenhoff2016), STD-LUCA [last common universal ancestor]/ Euk[Eukaryotes]/ Ver[Vertebrata]/ Fun[Fungi]; **(B)** Species Tree Discordance Benchmark (Altenhoff2009); columns distinguish realm (TT-Euk [Eukaryotes]/Fun[Fungi]); **(C)** Agreement with Reference Gene Phylogenies: SwissTree (Boeckmann 2011); **(D)** Agreement with Reference Gene Phylogenies: TreeFamA (Li 2006); **(E)** boxplot for true positive rate in **(F)**; (F) boxplot for positive predictive value rate in (C); **(G)** Gene Ontology conservation test; **(H)** Enzyme Classification (EC) conservation test.

## Supplementary Information

### Mapping of proteomes between QFO benchmark and eggNOG

Using the NCBI taxonomy table (ftp://ftp.ncbi.nih.gov/pub/taxonomy/taxdump.tar.gz) we matched the species by their taxonomy identifier, official name or by best suitable alternative (see TABLE). Subsequently we ran DIAMOND *blastp* [REF] to match individual proteins by reciprocal best hit (matching ratio median 0.98; max 1.0; min 0.78; 1.5IQR 0.88).

#### Conversion algorithm for orthologous groups to pairwise orthologs

To obtain pairwise orthologs for a pair of species in the QFO benchmark we (1) select the best fitting taxonomy level offered by eggNOG (e.g. for human and mouse it is superprimates/spriNOG), (2) filter the OGs of the best fitting level to the once containing both species, (3) generate all possible pairs between the genes of the two species in each OG. This simple procedure relies on the assumption that eggNOG has sufficient resolution in terms of taxonomic level for each species pair, such that the proteins in the relative OGs are indeed orthologs. Because this is not always possible we are aware of the method’s limitations and consider the optimization of this problem beyond the scope of this paper.

Table of manually selected mappings between QFO species and eggNOG specie

Table of manually selected mappings between QFO species and eggNOG species
| qfo_id | qfo_name | eggNOG_id | eggNOG_name | ncbi_id |
| --- | --- | --- | --- | --- |
| 5207 | cryptococcus neoformans | 283643 | cryptococcus neoformans var. neoformans b-3501a | 5207 |
| 83333 | escherichia coli | 511145 | escherichia coli str. k-12 substr. mg1655 | 83333 |
| 173 | leptospira interrogans | 267671 | leptospira interrogans serovar copenhageni str. fiocruz l1-130 | 173 |
| 6945 | ixodes scapularis | 6945 | ixodes scapularis | 6945 |
| 76856 | fusobacterium nucleatum | 190304 | fusobacterium nucleatum subsp. nucleatum atcc 25586 | 76856 |
| 478009 | halobacterium salinarum | 64091 | halobacterium sp. nrc-1 | 478009 |
| 184922 | giardia intestinalis | 184922 | giardia lamblia atcc 50803 | 184922 |
| 5476 | candida albicans | 573826 | candida dubliniensis cd36 | 5476 |
| 1773 | mycobacterium tuberculosis | 83332 | mycobacterium tuberculosis h37rv | 1773 |
| 375 | bradyrhizobium japonicum | 1037409 | bradyrhizobium japonicum usda 6 | 375 |

